# Resource: A Curated Database of Brain-Related Functional Gene Sets (Brain.GMT)

**DOI:** 10.1101/2024.04.05.588301

**Authors:** Megan H. Hagenauer, Yusra Sannah, Elaine K. Hebda-Bauer, Cosette Rhoads, Angela M. O’Connor, Stanley J. Watson, Huda Akil

## Abstract

Transcriptional profiling has become a common tool for investigating the nervous system. During analysis, differential expression results are often compared to functional ontology databases, which contain curated gene sets representing well-studied pathways. This dependence can cause neuroscience studies to be interpreted in terms of functional pathways documented in better studied tissues (*e*.*g*., liver) and topics (*e*.*g*., cancer), and systematically emphasizes well-studied genes, leaving other findings in the obscurity of the brain “ignorome”.
To address this issue, we compiled a curated database of **918** gene sets related to nervous system function, tissue, and cell types (“Brain.GMT”) that can be used within common analysis pipelines (*GSEA, limma, edgeR*) to interpret results from three species (rat, mouse, human). Brain.GMT includes brain-related gene sets curated from the Molecular Signatures Database (MSigDB) and extracted from public databases (GeneWeaver, Gemma, DropViz, BrainInABlender, HippoSeq) and published studies containing differential expression results.
Although Brain.GMT is still undergoing development and currently only represents a fraction of available brain gene sets, “brain ignorome” genes are already better represented than in traditional Gene Ontology databases. Moreover, Brain.GMT substantially improves the quantity and quality of gene sets identified as enriched with differential expression in neuroscience studies, enhancing interpretation.

**Graphical abstract:** 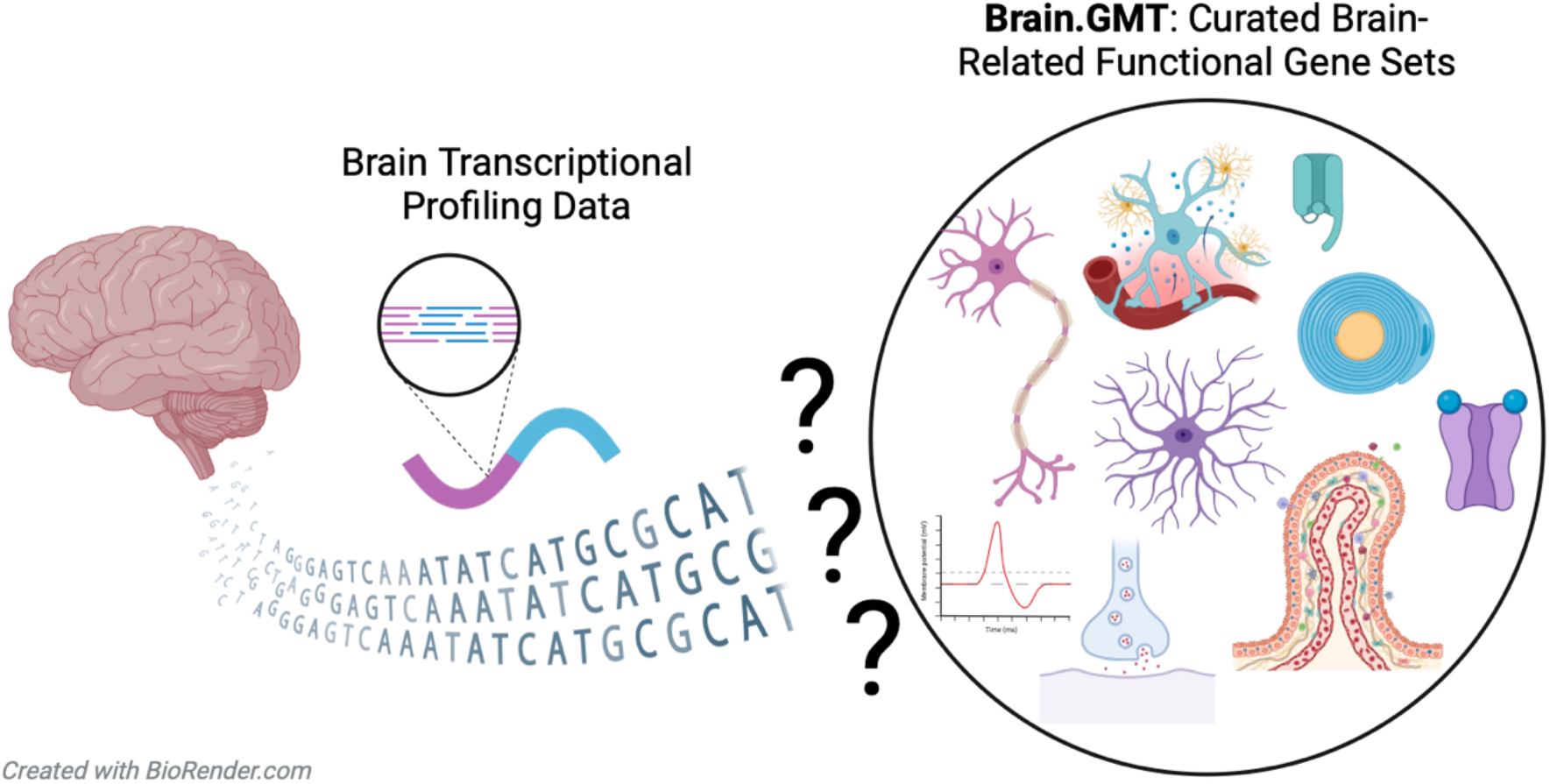

**Specifications table:** 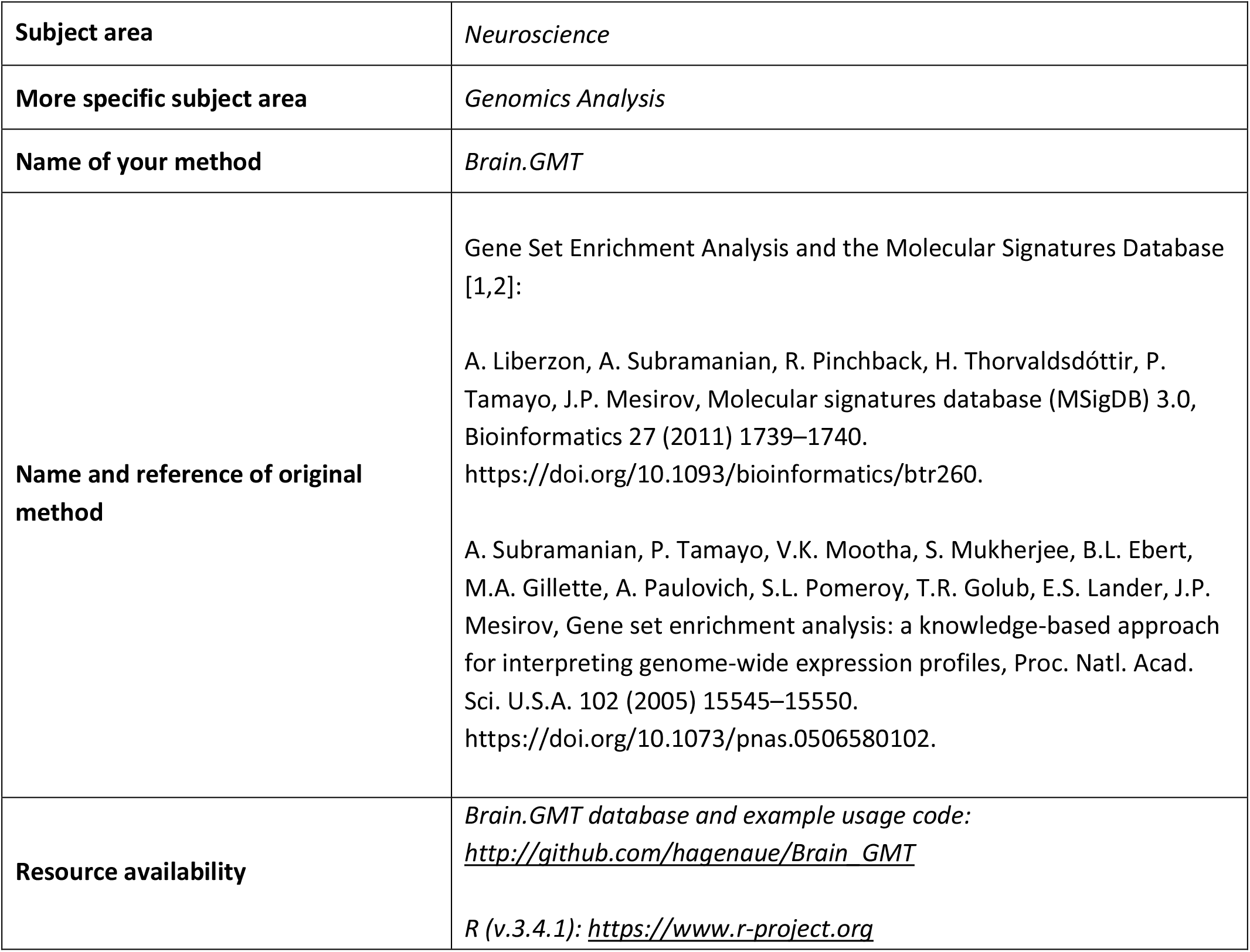

## Background

Over the past two decades, neuroscientists have embraced the use of transcriptional profiling technologies such as microarray and RNA-Sequencing (RNA-Seq). These technologies measure the expression of thousands of genes (transcripts) in each biological sample, providing a broad overview of cellular or tissue function. Using these technologies, neuroscientists can move beyond “hypothesis-driven” science - defined by preconceived notions of how the brain should function - and into the realm of unbiased discovery.

However, it can be challenging to interpret the differential expression results from transcriptional profiling studies. Often, researchers begin to assign biological meaning to differentially expressed genes by referencing large gene ontology or functional annotation databases that represent a curation of consolidated knowledge from published literature (*e*.*g*., Gene Ontology Consortium [3], Kyoto Encyclopedia of Genes and Genomes [4], Reactome [5]). Many tools are available for formally comparing differential expression results to gene ontology databases (e.g., GORilla [6], DAVID [7], EnrichR [8]). These tools typically determine whether groups of genes representing particular functional pathways or biological processes (gene sets) show a significant enrichment of differential expression within the results – *i*.*e*., more differential expression than expected by random chance. Within R analysis pipelines, common algorithms for conducting these analyses (e.g., Gene Set Enrichment Analysis (GSEA) [2], CAMERA [9], ROAST [10], ROMER [11]) use gene set database files in the Gene Matrix Transposed format (.gmt) available at the Molecular Signatures Database (MSigDB [1]) and elsewhere [8].

Like many neuroscientists, we have found that comparing our brain-derived differential expression results to traditional gene ontology databases is often unenlightening. Many gene sets in these databases are derived from better studied tissues (*e*.*g*., liver) and topics (*e*.*g*., cancer), with questionable relevance to brain function (e.g., “SPERM MOTILITY”, “HEART MORPHOGENESIS”).

Moreover, the use of gene ontology databases for two decades to interpret differential expression results has caused a “bandwagon effect”, encouraging the promotion of well-studied genes in discussions and abstracts. One study estimated that just 5% of brain-expressed transcripts were the focus of 70% of the neuroscience literature, and 20% had almost no representation at all – a subset referred to as the “brain ignorome” [12].

To improve the interpretation of brain-derived differential expression results, we compiled a custom gene set database (Brain.GMT) focused on sets of genes associated with brain function, brain cell-types, brain co-expression networks, and regional gene expression signatures. We initially constructed Brain.GMT as part of projects using hippocampal [13–16] and nucleus accumbens tissue [15] from rodent models of neuropsychiatric disorders. To rapidly compare our results to existing literature, we also constructed gene sets using differential expression results from related publications and differential expression databases.

This paper serves as detailed methodological documentation to accompany our transcriptional profiling studies using Brain.GMT [13–16]. Since we have found Brain.GMT to be exceptionally useful, we also provide detailed guidance to accompany its public release for use by other researchers. Finally, Brain.GMT can serve as a case study demonstrating the utility of customized gene set databases for the interpretation of differential expression results, guiding future development efforts.

## Method details

### General Methods

#### Overview of the .GMT gene set database format

The Gene Matrix Transposed file format (*.gmt) is used to input a database of gene sets into genomics analysis pipelines like Gene Set Enrichment Analysis (GSEA: [2]), Limma [11], and edgeR [17]. This file format is a tab delimited text file, with each row representing a particular gene set. The first column includes the name/identifier for the gene set (string: free text), the second column contains information regarding the source of the gene set (string: free text), and then there are columns listing each of the genes included in the gene set (one gene identifier per cell). Traditionally, the annotation used for the listed genes is official gene symbol, with a key .gmt provider, the Molecular Signatures Database (MSigDB: *http://software.broadinstitute.org/gsea/msigdb/index.jsp*, [1,2]), focusing on human gene symbols and orthologs. Since we initiated our project, MsigDB has also begun providing .gmt files focused on mouse symbols and orthologs [18], but those resources were not available at the time that we were conducting our work. Our laboratory analyzes differential expression results from three species (rat, mouse, human), so we constructed three versions of Brain.GMT that list the gene set constituents using official gene symbols from rats, mice, and humans, respectively.

#### General Methods Used for Gene Set Construction

For all custom gene sets, gene symbol annotation was obtained from the original study/database or translated from the gene annotation provided by the source material (e.g., Ensembl ID, Entrez ID) into official gene symbol using relevant annotation packages (org.Hs.egSymbol v.3.4.1 [19], org.Mm.egSymbol v.3.4.1 [20], org.Rn.egSymbol v.3.4.1 [21]). Only unique gene symbols were included in the final gene set (no duplicates). If the gene symbols provided by the original study/database included older, date-related gene symbols that cause problems when imported into Microsoft Excel (*March* genes, *Sept* genes, *Dec* genes, *Nov*), they were changed to updated nomenclature. Then, when appropriate, species (rat, mouse, human) orthologs for the genes included in each gene set were identified using the ortholog database on the Mouse Genome Initiative (MGI) website [22] (*http://www.informatics.jax.org/homology.shtml*, downloaded 02/28/2021).

While constructing gene sets from the various source material, we targeted a gene set size that would be easily compatible with common analysis pipelines (gene set sizes ranging from 10-999 genes), using database or publication-specific statistical thresholds to define the genes included in each gene set. When possible, we included separate gene sets for genes that were upregulated and downregulated within a particular condition.

#### Database and Code Availability

The most recent version of Brain.GMT (v.2) is available on our Github site for the analysis of genomics results from rats, mice, and humans (*http://github.com/hagenaue/Brain_GMT*).

The R code used to construct Brain.GMT has been released on our Github site (Rstudio v.1.0.153, R v.3.4.1, *https://github.com/hagenaue/Brain_GMT/tree/main/Code*). We have also provided example R code illustrating the use of Brain.GMT within a Fast Gene Set Enrichment Analysis (fGSEA, [23]): (*https://github.com/hagenaue/Brain_GMT/blob/main/BrainGMT_exampleUsage.R*).

### Methods for Gene Set Construction

#### Overview of Included Gene Sets

Our custom gene set file (Brain.GMT: 918 gene sets, **Table 1**) was designed to provide greater insight into brain-derived differential expression results than traditional functional ontology. Originally, the gene set file was constructed as part of projects using rodent models of neuropsychiatric disorders performed on tissue from the hippocampus [13–16] and nucleus accumbens [15], and thus emphasizes gene sets derived from related regions and topics.

**Table 1.** An Overview of the Gene Sets Included in Brain.GMT. The Brain.GMT project was originally initiated to provide insight into hippocampal differential expression (DE) studies related to neuropsychiatric disorder (v.1), and then expanded to include gene sets specific to the nucleus accumbens (v.2). The source for each variety of gene set is referenced above, along with a brief description of the type of gene set included, and tissue. Also noted is whether the gene sets were extracted from the source following additional curation by a trained neuroscientist for relevance to the nervous system or project themes, and the final number of gene sets included from the source in Brain.GMT.

| Brain.GMT Gene Sets: |  |  |  |  |  |
| --- | --- | --- | --- | --- | --- |
| Version of Brain.GMT | Source | Type of Gene Set | Tissue Source | Curated for Brain Relevance | # of Gene Sets Included in Brain.GMT |
| 1 | MSigDB: "C2: Curated Gene Sets" (Liberzon et al. 2011) | Curated Gene Sets | Nervous System | Y | 158 |
| 1 | MSigDB: "C8: Cell Type Signature Gene Sets" (Liberzon et al. 2011) | Cell Type Enriched Expression | Nervous System & Blood | Y | 211 |
| 1 | BrainInABlender (Hagenauer et al. 2018) | Cell Type Enriched Expression | Nervous System (especially Cortex) | N | 39 |
| 1 | DropViz (Saunders et al. 2018) | Cell Type Enriched Expression | Hippocampus | N | 13 |
| 2 | DropViz (Saunders et al. 2018) | Cell Type Enriched Expression | Nucleus Accumbens | N | 12 |
| 1 | HippoSeq (Cembrowski et al., 2016) | Regional Enriched Expression | Hippocampus | N | 14 |
| 1 | Coexpression Analyses: (Johnson et al. 2015, Park et al. 2011) | Coexpression Networks | Hippocampus | N | 55 |
| 1 | Curated in (Birt et al. 2021) | Published DE Results: Selective Breeding for Internalizing Behavior | Hippocampus | N | 19 |
| 1 | Hope For Depression Research Foundation: (Gray et al. 2014; Bagot et al. 2016, Bagot et al. 2017, Pena et al. 2019) | Published DE Results: Stress Interventions | Hippocampus | N | 14 |
| 1 | Meta-Analyses: (Gandal et al. 2018) | Published DE Results: Neuropsychiatric Disorder Meta-Analyses | Cortex | N | 14 |
| 2 | GeneWeaver (Baker et al. 2012) | Published DE Results: Stress, environmental enrichment, affective behavior, and mood disorder | Nucleus Accumbens | Y | 6 |
| 1 | GeneWeaver (Baker et al. 2012) | Published DE Results: Stress, environmental enrichment, affective behavior, and mood disorder | Hippocampus | Y | 33 |
| 2 | Gemma (Zoubariev et al. 2012) | DE Reanalysis Pipeline: Stress, environmental enrichment, affective behavior, and mood disorder | Nucleus Accumbens | Y | 29 |
| 1 | Gemma (Zoubariev et al. 2012) | DE Reanalysis Pipeline: Stress, environmental enrichment, affective behavior, and mood disorder | Hippocampus | Y | 301 |
| **Total** |  |  |  |  | **918** |
| **Packaged with Traditional Ontology:** |  |  |  |  |  |
| 1 | MSigDB: "C5: GO Gene Sets" (Liberzon et al. 2011) | Traditional Gene Ontology | Generic | N | 14,996 |

To provide insight into how to interpret our differential expression results in terms of brain function, broadly speaking, we included brain-related functional gene sets and brain cell-type related gene sets that were scraped from the Molecular Signatures Database [1,2], BrainInABlender [24], and DropViz [25], as well as a few gene sets related to brain co-expression networks and regional gene expression signatures [26–28].

To this file, we added additional gene sets specifically designed to provide insight into the role of the hippocampus and nucleus accumbens in processing affective behavior. We started by creating gene sets that would allow us to quickly and uniformly assess the overlap of our results with the findings from related publications, including the effects of stress in the hippocampus and nucleus accumbens identified by other members of the Hope for Depression Research Foundation [29–32], the effects of selective breeding targeting internalizing-like behavior in the hippocampus (curated in [13]), and the effects of human neuropsychiatric disorders as documented within some of the largest differential expression meta-analyses available at the time (cortex: [33]). Then, to gain a more comprehensive comparison, we extracted the differential expression results from all studies in the hippocampus and nucleus accumbens related to stress, enrichment, social and affective behavior, and mood disorder in two online databases of differential expression results: Gemma [34,35] and GeneWeaver (*https://www.geneweaver.org*, [36]).

To create a more well-rounded picture, we packaged Brain.GMT with a traditional commonly-used collection of gene ontology gene sets included in the Molecular Signatures Database [1,2] (MSigDB v7.3, *http://software.broadinstitute.org/gsea/msigdb/index.jsp*, downloaded 2021-03-25) (“C5: GO Gene Sets”, file: “c5.all.v7.3.symbols.gmt.txt”, # of gene sets: 14,996).

### Detailed Methods for Constructing Database Derived Gene Sets

#### MSigDB-Derived Brain-Related Gene Sets

Within the Molecular Signatures Database (MSigDB, *http://software.broadinstitute.org/gsea/msigdb/index.jsp*, [1,2]) there are two commonly used gene set collections that include several hundred brain-related gene sets (“C2: Curated Gene Sets”, “C8: Cell Type Signature Gene Sets”). We downloaded these gene set collections (MSigDB v7.3, downloaded 2021-03-25: files: “c2.all.v7.3.symbols.gmt.txt”, “c8.all.v7.3.symbols.gmt.txt”) and a trained neuroscientist curated and filtered them for specific relevance to nervous system tissue and function, including gene sets related to nervous system cell types and blood cell types (as blood is often present in nervous system tissue), neurological disorders, psychiatric disorders, neurotransmission, psychoactive drugs, neuroactive hormones, stress response, and gene sets derived from a variety of other studies conducted using central nervous system tissue (“C2: Curated Gene Sets”: # of filtered gene sets: 158; “C8: Cell Type Signature Gene Sets”: # of filtered gene sets: 211).

#### DropViz-Derived Gene Sets Related to Brain Cell Types

DropViz is a database of single cell RNA-Seq (scRNA-Seq) results from central nervous system tissues [25]. To gain better insight into differential expression related to the cell types present in our brain regions of interest, we extracted brain cell-type enriched gene sets from the DropViz database using the results from hippocampal and nucleus accumbens tissue (*http://dropviz.org*, accessed March 25, 2021). We extracted the results for genes that had enriched expression in each of the cell types (Cell Type Cluster vs. Rest of Region: p-value< 10^−30^, minimum fold ratio=4); a greater level of specificity was difficult to achieve for many neuronal subtypes. To reduce noise, we required minimum expression levels within the cell type of interest (minimum logCPM in Cell Type Cluster=0.5) and excluded genes that were also strongly expressed in the rest of the tissue (maximum expression levels logCPM in Rest of Region=6). When possible, to improve specificity, the gene sets associated with the cell type clusters from the DropViz database were further filtered to include either 1) all genes with fold change greater than 10 for the cluster vs. the rest of the brain region (if there were more than 50 genes meeting these criteria), or 2) The top 50 genes with the highest fold change for the cluster vs. the rest of the region (# of gene sets: 25).

#### GeneWeaver-Derived Gene Sets

GeneWeaver is a web-based curated repository of genomic experimental results with accompanying toolsets [36]. With the help of the developer, Dr. Erich Baker, we extracted public experimentally-derived gene sets from the GeneWeaver database (*https://www.geneweaver.org,* accessed June 28 2021) for studies from the nucleus accumbens or hippocampus related to stress, environmental enrichment, affective behavior, and mood disorder. The results were ranked by the differential expression metric provided (false discovery rate (FDR), p-value or absolute effect size), and the gene symbol annotation for the top 25 results (or full results, if <25) was extracted, ignoring results lacking gene symbol annotation or mapped to multiple gene symbols (# of gene sets: 38).

#### Gemma-Derived Gene Sets

Gemma is a large web database of curated and re-analyzed gene expression studies [34,35]. We extracted experimentally-derived gene sets from the Gemma database (*https://gemma.msl.ubc.ca/home.html*) using the gemmaAPI (Github: PavlidisLab/gemmaAPI.R) to access differential expression results. We used *annotationInfo()* to download a list of all datasets including the annotation “nucleus accumbens” or “hippocampus” (nucleus accumbens: accessed June 3, 2021, hippocampus: accessed June 15, 2021), and narrowed that list to public datasets from humans, mice, or rats that weren’t tagged as troubled (nucleus accumbens: 103 datasets, hippocampus: 648 datasets). Datasets that were tagged as having batch confounds were reviewed by hand to ascertain whether the confound would interfere with the interpretation of the variable of interest. Datasets were then further reviewed by hand for relevance to stress, environmental enrichment, affective behavior, and mood disorder (NACC: 15 datasets, HC: 86 datasets).

The results for the datasets of interest were then downloaded locally (accessed June 24, 2021).

The “analysis.results.txt” file for each dataset, which included the p-values and q-values for each variable in the dataset for each transcript/gene, was extracted and joined with the “resultset” for each variable, which included the FoldChange, T-stat, and P-value outputted for each contrast, using the database unique gene identifier (“Element_Name”). These results were then filtered to remove results that either lacked gene symbol annotation or that had mapped to multiple gene symbols (separated by a “|” in the database). To produce gene sets of the targeted size (10-999 genes), these files were subsetted to pull out results for each variable that survived a threshold of false discovery rate (FDR)<0.10 and p-value<0.0001, and then the results for the specific contrasts for that variable were further filtered using p<0.05. The down-regulated (FoldChange<1) and up-regulated (FoldChange>1) results were divided into separate gene sets. These gene sets were then ranked by FoldChange, and only the 999 most down-regulated and 999 most up-regulated transcripts were maintained in the gene set. The final database included 329 gene sets (NACC: 29, HC: 301).

### Detailed Methods for Constructing Publication Derived Gene Sets

#### Co-expression Networks

We added a set of custom gene sets that had been previously curated [37] to summarize hippocampal co-expression networks [27,28] (# of gene sets: 55).

#### Regional and Cell-Type Enriched Expression

We added a set of custom gene sets that had been previously curated [37] to summarize hippocampal regional gene expression signatures (HippoSeq: [26], # of gene sets: 14) and gene sets enriched for expression within specific brain cell types (BrainInABlender database [24] (*https://github.com/hagenaue/BrainInABlender*, v.0.0.0.9000, # of gene sets: 39)

#### Stress and Psychiatric Disorder-Related Gene Sets

We also created gene sets that would allow us to quickly assess the overlap of our differential expression results with the findings from related publications. We started by including gene sets representing the stress-related differential expression identified in the hippocampus and nucleus accumbens by other members of our research consortium (the Hope for Depression Research Foundation). This included gene sets derived from chronic restraint stress, forced swim stress, and acute corticosterone in the hippocampus ([31]: Suppl. Tables 2, 3, 6, 7) which we filtered to produce gene sets within the targeted size range (10-999 genes) using p<0.005 for any of the individual comparisons, divided into upregulated and down-regulated for each comparison, or p<0.00005 for an ANOVA encompassing all conditions. This also included gene sets related to chronic social defeat stress in the hippocampus or nucleus accumbens (filtered to p<0.005 in addition to using publication-defined thresholds: ([29]: Table S1: p<0.05, |FC|>1.3), ([30]: Table S2, S4: p<0.05, |FC|>1.3), [32]: Suppl Data 2: |FC|>30%) (# of gene sets: 14).

**Table 2.**
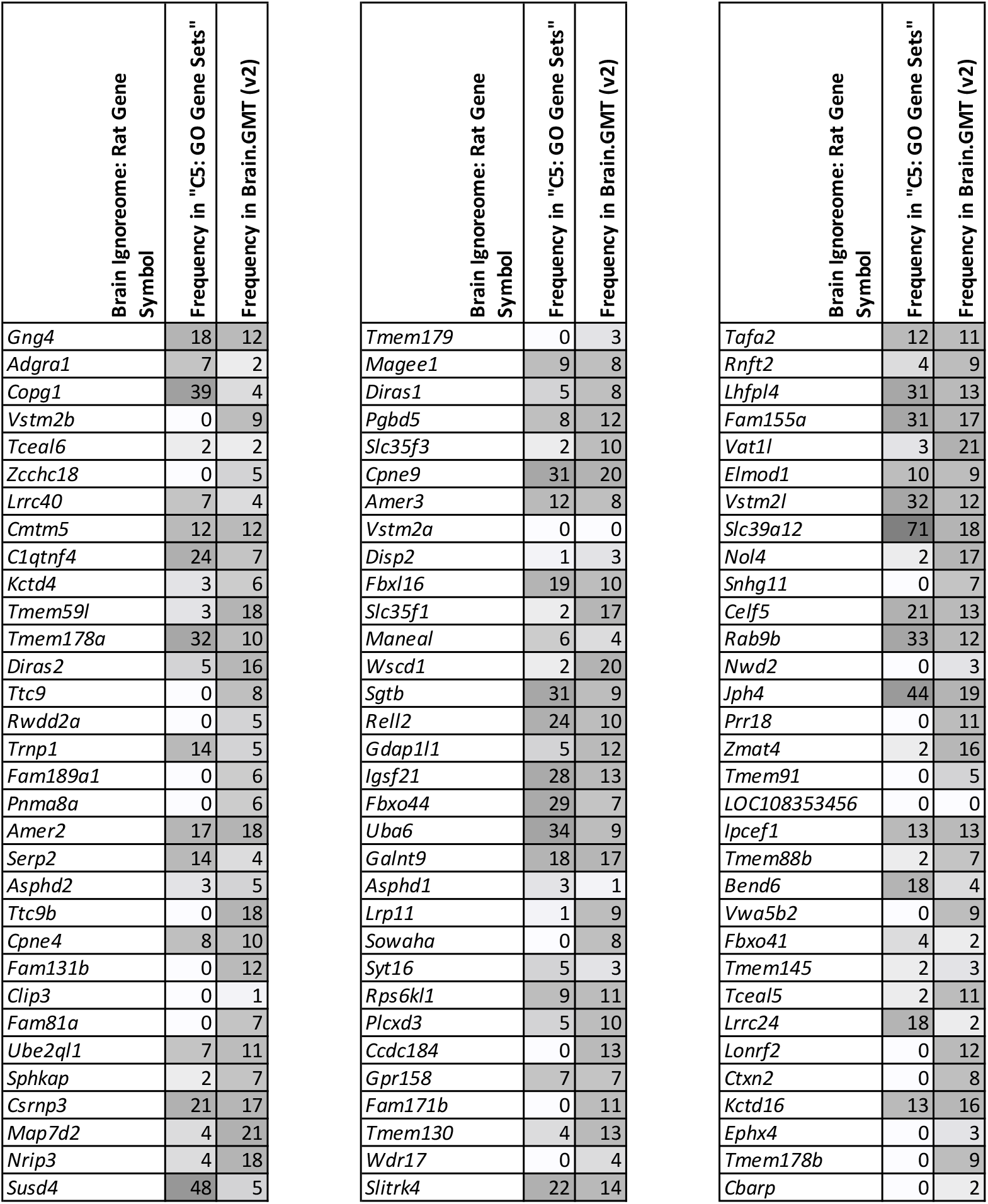
“Brain ignorome” genes are better represented in Brain.GMT than in traditional Gene Ontology. The table shows the frequency that the “Brain Ignorome” genes identified in [12] show up in a traditional gene ontology database (MSigDB’s “C5: GO gene sets”: 14,996 gene sets; rat orthologs) in comparison to Brain.GMT (rat, v.2, 918 gene sets). Grey scale is used to make frequency values easier to visualize (white= lowest frequency, dark grey=highest frequency). The order of the gene symbols follows the original supplementary table in [12]. The table is split into three for the purpose of fitting easily on a page.

We added gene sets from hippocampal transcriptional profiling studies examining the effects of selective breeding targeting internalizing behavior [37–45]. These differentially expressed gene lists had been curated in a previous publication [37] using their original publication-specific criteria to define significance. We created up-regulated and down-regulated versions of each gene set when there was a sufficient number of differentially expressed genes (>10) (# of gene sets: 19).

Finally, we compiled a set of gene sets related to human neuropsychiatric disorders (Major Depressive Disorder, Bipolar Disorder, Schizophrenia, Autism Spectrum Disorder, Alcohol Abuse Disorder) using the differentially expressed genes identified in one of the largest meta-analyses of brain transcriptional profiling studies conducted at that time (using cortical tissue, [33]: filtered to produce gene sets within the targeted size range (10-999 genes) using FDR<0.05 & p<0.001). Each of these gene sets was divided into down-regulated and upregulated genes (# of gene sets: 14).

### Methods for Demonstrating Utility

#### The Representation of “Brain Ignorome” Genes in Brain.GMT

To demonstrate the potential for Brain.GMT to improve the interpretation of brain-related genomics results, we compared the representation of “brain ignorome” genes (all genes listed in Table S5 of [12]) in Brain.GMT (v.2., # of gene sets: 918) as compared to a traditional functional ontology database (the MSigDB “C5: GO Gene Sets”, packaged with Brain.GMT, # of gene sets: 14,996). Due to the focus of our laboratory’s current projects, we chose to run this comparison using the rat version of Brain.GMT (packaged with the rat orthologs for MSigDB’s “GO Gene Sets”) and the rat orthologs for the “brain ignorome” genes (orthologs determined using RGD.mcw.edu, accessed 05-22-2023).

#### Trial Runs Using Brain.GMT within Gene Set Enrichment Analyses

To illustrate the benefits of using Brain.GMT within gene set enrichment analyses of brain differential expression results, we referenced the results from three previous publications that trialed our gene set database [14–16]. Each of these studies focused on rodent (rat, mouse) models of mood disorder, behavioral temperament, or stress response using tissue from the hippocampus or nucleus accumbens. These samples represented both sexes, with a skew towards males: the results from [14] reflected a sample evenly composed of males and females, with a similar relationship between gene expression and internalizing behavior observed in both sexes, whereas the results from [15,16] reflected all male samples. In each study, we used a .GMT file containing both the Brain.GMT gene sets and traditional gene ontology gene sets (**Table 1**) as input while conducting a Fast Gene Set Enrichment Analysis (fGSEA, [23]) of our differential expression results (versions for each publication: [14,16]: rat Brain.GMT v.1, [15]: rat Brain.GMT v.2).

For each of these studies, the analysis methods, code, inputted differential expression results, and outputted gene set enrichment results were released as part of their respective publications. For [14], the referenced fGSEA results are from worksheet 2 (“Directional_Test”) in **Supplemental Table S5**. For [15], the referenced fGSEA results are from worksheets 2 and 3 (“SortbyEE” and “SortbySD”) from both **Tables S4 and S5**, with the false discovery rate defined by the minimum FDR from the Model 1 and Model 2 analyses (“EE_Min_AdjPval” and “SD_MIN_AdjPval). For [16], the referenced fGSEA results are from the code release accompanying the publication (*https://github.com/hagenaue/HDRF_MetaAnalysis_Downstream*).

We also trialed the use of Brain.GMT (v.2) as part of a fGSEA analysis performed on differential expression results from a meta-analysis of the effects of sleep deprivation in the cortex in rodents (rats/mice) as measured by microarray or RNA-Seq. Since this work is unpublished, we have only briefly called out our findings as a point of comparison to the studies from the hippocampus and nucleus accumbens.

### Method Validation

#### “Brain Ignorome” genes are better represented in the gene sets in Brain.GMT

We have found that Brain.GMT greatly improves the interpretation of brain differential expression results, even though the database is still undergoing development and currently only represents a fraction of available brain gene sets. The ‘brain ignorome” genes [12] are already better represented in Brain.GMT than in traditional Gene Ontology. For example, 28% of the “brain ignorome” genes (27 of 96) had no representation in MSigDB’s traditional gene ontology collection (“C5: GO gene sets”: 14,996 gene sets) but only 2% (2 of 96) lacked representation in Brain.GMT (v.2, 918 gene sets) (**Table 2**). Moreover, even though Brain.GMT (v.2) currently only contains 918 gene sets, the “brain ignorome” genes were represented in a median of nine Brain.GMT gene sets apiece (range: 0-21, average: 9.5) but only in a median of four of MSigDB’s traditional gene ontology gene sets (“C5: GO gene sets”) (range: 0-71, average: 10.3). Considering the number of gene sets included in each collection, the “brain ignorome” genes were on average more than 15X more likely to show up in any particular gene set within Brain.GMT compared to within MSigDB’s “C5: GO gene sets”.

#### Overview of trial runs using Brain.GMT for Gene Set Enrichment Analysis performed on brain-derived differential expression results

We have now used the Brain.GMT custom gene set database to improve our interpretation of differential expression results within three publications [14–16] and one unpublished study (Rhoads et al., *in preparation*). Each of these studies focused on rodent (rat, mouse) models of mood disorder, behavioral temperament, or stress response. In each study, we used a .GMT file containing both the Brain.GMT gene sets and traditional gene ontology gene sets (**Table 1**) as input while conducting a Gene Set Enrichment Analysis (fGSEA, [23]) of our differential expression results. In the case of meta-analyses ([16], Rhoads et al. *in preparation*), we removed any gene sets that referenced datasets included in our meta-analysis.

#### Gene sets in Brain.GMT were more likely to be enriched with brain differential expression

In each case, we found that a disproportionate number of the gene sets detected as being significantly enriched with differential expression (FDR<0.05) came from the Brain.GMT gene set database and not traditional ontology (**Figure 1**). Within our gene set enrichment results, a large percent of the gene sets that were significantly enriched with differential expression (FDR<0.05) were from Brain.GMT (vs. traditional gene ontology), ranging from 26% to 61%. In contrast, within the full gene set enrichment results (both significant (FDR<0.05) and not significant (FDR>0.05)), the percent of gene sets that were from Brain.GMT (vs. traditional gene ontology) was only around 7%. This disproportionate representation was even more evident within the strongest results – when ranked by p-value or normalized enrichment score, it was not uncommon for almost all of the top 10 results to be gene sets from Brain.GMT.

**Figure 1.**
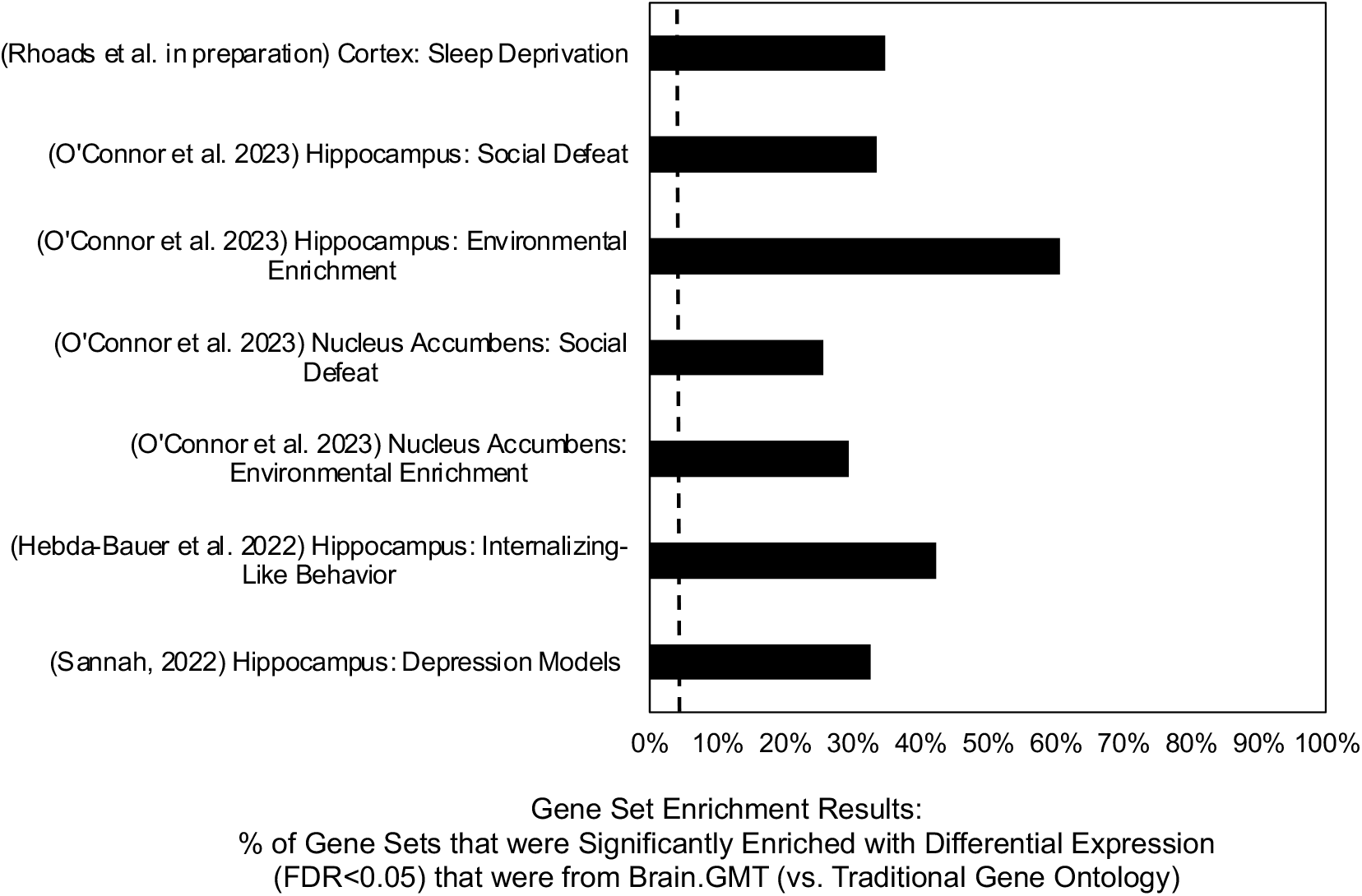
Gene Set Enrichment Analysis of brain differential expression results using. **a .GMT containing both Brain.GMT and traditional Gene Ontology gene sets shows disproportionate enrichment in Brain.GMT gene sets.** In each study, we used a .GMT file containing both the Brain.GMT gene sets and traditional gene ontology gene sets (**Table 1**) as input while conducting a Gene Set Enrichment Analysis (fGSEA, [23]) of our differential expression results. The number of gene sets included in the final results varied by study based on dataset characteristics and fGSEA filtering parameters, but in all cases the percent of gene sets that were from Brain.GMT in the full results (vs. traditional gene ontology) hovered around 7% (dashed line). In contrast, the percent of the gene sets that were significantly enriched for differential expression (FDR<0.05) that were from Brain.GMT (vs. traditional gene ontology) was much higher, ranging from 26% to 61% (black bars).

#### The gene sets from Brain.GMT that were enriched with differential expression were easier to interpret

The Brain.GMT gene sets also improved the interpretability of the differential expression results. This was particularly striking within our meta-analysis of gene expression in the hippocampus across animal models of depression [16], where the strongest pattern in the results was down-regulation within Brain.GMT gene sets representing glial-enriched expression, particularly astrocytes, in a manner paralleling previous findings in depressed human patients (e.g., [46]). The Brain.GMT gene sets also helped disambiguate the enrichment of differential expression within traditional gene ontology gene sets. For example, significant enrichment of differential expression within the gene ontology gene sets of GOBP_HEART_MORPHOGENESIS, GOBP_RENAL_SYSTEM_VASCULATURE_DEVELOPMENT, and GOBP_MITRAL_VALVE_DEVELOPMENT were much easier to interpret when accompanied by a stronger enrichment of differential expression within a variety of Brain.GMT gene sets representing brain endothelial cell and brain mural cell-related gene expression [15], or when we observed significant enrichment within the gene ontology gene sets of GOBP_INNATE_IMMUNE_RESPONSE and GOBP_DEFENSE_RESPONSE_TO_OTHER_ORGANISM it was useful to know that there was also stronger enrichment within a variety of Brain.GMT gene sets representing microglial-related gene expression (brain immune cells) [14]. Likewise, the enrichment of differential expression results within gene ontology gene sets like GOCC_CILIUM, GOBP_SPERM_MOTILITY, and HP_MALE_INFERTILITY seemed completely incomprehensible until we had the added context of much stronger differential expression within the Brain.GMT gene sets containing ependymal cell markers (ciliated brain cells) [15].

#### A custom gene set database (Brain.GMT) was useful for making formal, pre-specified comparisons with published literature

We have found that the ability to include gene sets within Brain.GMT that allow us to rapidly compare our differential expression results to previous differential expression studies on related topics has also been a boon. Within our study examining the effects of selective-breeding and genetic propensity for internalizing behavior on hippocampal gene expression, we found that our results showed a strong enrichment of differential expression within sets of genes identified as differentially expressed in the hippocampus of a related, independent rodent model [14], allowing us to feel more confident that our results were broadly generalizable and not an artifact of genetic drift within our colony. Within our study examining the effects of adolescent exposure to environmental enrichment and social defeat, we found an enrichment of differential expression within a disproportionate percent of gene sets related to our interventions and affected behaviors, including aggression, social behavior, and activity levels [15]. Moreover, the use of a formalized gene set enrichment analysis forced us to conduct comparisons between our findings and previous publications in a more comprehensive, standardized way that included a multiple comparisons correction for the number of comparisons made and required pre-specification of all desired comparisons, decreasing the temptation to cherry pick examples that supported our findings from the results sections of previous publications.

#### Limitations and Future Directions

We have started to regularly use Brain.GMT in our differential expression analyses because it has turned out to be incredibly beneficial for guiding interpretation. That said, Brain.GMT is still undergoing development and currently only represents a fraction of available brain gene sets. There are also notable limitations to its usage and interpretation that should be considered prior to use. It is, in many ways, more of a prototype that proves the benefits of further development than a finished product, but still represents a notable improvement over the status quo.

#### Considerations When Interpreting Brain.GMT Results

##### Bias in favor of coding genes

There are several important weaknesses to consider when interpreting results from Brain.GMT that are also typical of .gmts from other popular databases, such as MSigDB. One important weakness is the dependency on official Gene Symbols as the identifier for gene set constituents. Gene Symbols can be unstable gene identifiers, especially for genes that have been recently characterized. Moreover, not all genes have Gene Symbols, especially non-coding genes. This means that although Brain.GMT provides much better representation of the “brain ignorome”, the genes represented in the gene sets in Brain.GMT are still skewed in favor of better-studied, coding genes. When referencing gene sets that were originally derived in a different species, this bias is heightened due to the difficulty of identifying orthologs for non-coding genes. In the future, it would be useful to construct versions of Brain.GMT that use more stable and less biased identifiers, like Ensembl IDs.

##### Gene set definitions vary by source material

Another important consideration when interpreting results from Brain.GMT that are also typical of .gmts from other popular databases is the criteria for inclusion of a gene in a gene set varies based on the source material. For example, a gene set defined as including genes with astrocyte-enriched expression within BrainInABlender may use stricter cut-offs (e.g., 20-fold enrichment) than a gene set scraped from DropViz, or a gene set defined by differential expression in the hippocampus in the GeneWeaver database may use a different threshold for significance than a gene set scraped from the Gemma database. Therefore, if results from an analysis using Brain.GMT include an enrichment of differential expression within similar gene sets derived from one source and not another, this could reflect varying amounts of noise or specificity allowed by the original gene set definitions. Likewise, depending on the source material, a gene set may include all differentially expressed genes for a condition or may be divided into two gene sets representing upregulated and downregulated expression. If Brain.GMT is used within an analysis that considers the direction of effect of the differential expression results (e.g., fGSEA), there may be a bias against the gene sets that include all differential expression (both upregulated and downregulated) associated with conditions.

##### Overrepresentation of specific categories of gene sets

Likewise, when examining the top results from any gene set related analysis, including Brain.GMT, it is important to consider the prevalence of different types of gene sets within the .gmt database, as false positives will be more likely to reflect gene sets within prevalent categories. Within results using traditional ontology gene sets, this often leads to “cancer” related gene sets showing up amongst the top hits. Within Brain.GMT, or any gene set database customized to include more gene sets related to the tissue or topic of interest, these false positives may be harder to spot, as they are more likely to be believable results. For that reason, when using Brain.GMT, or other custom gene set databases, we recommend either using a stronger false discovery rate correction (e.g., FDR<0.01 instead of FDR<0.05) or taking into consideration the prevalence of various categories of gene sets when considering the enrichment results. For example, within one of our recent studies [15], we considered the percent of gene sets enriched with differential expression within particular pre-specified categories (e.g., mood disorder-related gene sets, stress-related gene sets) and highlighted categories with disproportionate enrichment in addition to examining the enrichment results for individual gene sets.

##### Shared artifacts and generic pathways driving overlap with differential expression results from previous studies

Finally, perhaps the most important consideration for interpreting the results from Brain.GMT – or from any direct comparison of differential expression findings – is the likelihood of observing an enrichment of differential expression within gene sets that are derived from differential expression studies that included similar, common sources of confounding variability. Transcriptional profiling studies are often weakly powered due to the expense of the methodology, making it impossible to reliably detect even moderately large effect sizes. As the biological effects of interest are often a magnitude smaller than highly-impactful technical artifacts, any slight imbalance in the experimental design can cause the top differential expression results to be mostly driven by technical factors such as dissection batches and variability in RNA quality. Therefore, an enrichment of brain-derived differential expression within a gene set derived from another differential expression study examining the effects of stress within brain tissue could imply that there are common mechanisms activated in the two studies, but it could also potentially imply that both studies shared a similar, common technical confound.

Moreover, some biological pathways are activated under a wide variety of conditions, such as the immediate early genes or inflammatory pathways [47], which can also drive an illusion of similarity when comparing the results from brain-derived differential expression studies.

Due to these issues, we found that enrichment of differential expression within Brain.GMT gene sets derived from weakly-powered individual differential expression studies (*e.g*., many of the gene sets scraped from smaller studies within GeneWeaver, Gemma, and individual publications) were harder to interpret than enrichment of differential expression within Brain.GMT gene sets derived from meta-analyses, higher powered studies, and studies characterizing large effects (e.g., cell type specific expression, effects of selective breeding). However, we also found that many of these issues with interpretation were easier to spot when using Brain.GMT within a formalized gene set enrichment analysis than when simply comparing differential expression results to the published literature or directly to the results of individual studies. Because many of the gene sets within Brain.GMT were divided into two gene sets representing upregulated and downregulated expression in relationship with the variables of interest, and Brain.GMT includes gene sets from differential expression results from a variety of related studies, it is easy to red flag results that show a pattern of enrichment within gene sets reflecting contradictory effects, and then examine the lists of leading genes for evidence of influential artifacts. For example, within one of our recent studies [15] we were excited to see that our stress-related differential expression results showed an enrichment within many gene sets related to fear conditioning. However, upon closer examination, we discovered that many of these findings indicated a contradictory direction of effect, and the leading genes driving the enrichment of differential expression in these gene sets were often immediate early genes, like *Fos* or *JunB*, which are highly reactive in the brain under a wide variety of conditions.

#### Expanding and Customizing Brain.GMT Gene Sets

There are many gene sets that could be added to Brain.GMT to increase functionality or tailor the database to the needs of other projects. For example, when scraping gene sets from Gemma, GeneWeaver, and Dropviz, we specifically focused on gene sets that would help provide needed insight into our current projects. Depending on the needs of future projects, it would be helpful to adapt our current code to extract gene sets from other central nervous system tissues or related to other research themes. There are also a variety of other useful types of brain-related gene sets that could be added with some additional effort. For example, Enrichr [8,48,49] includes a variety of downloadable gene set libraries (*https://maayanlab.cloud/Enrichr/#libraries*). These include some gene set libraries that are already centered on themes related to the central nervous system (*e.g*., Allen Brain Atlas identified cell types), and many libraries that are likely to include some gene sets derived from central nervous system tissue or related to central nervous system functions (*e.g*., gene sets implicated in neurological and behavioral phenotypes by Mouse Genome Informatics).

##### Reducing redundancy in gene set content

When adding or replacing gene sets in Brain.GMT, one important consideration is redundancy. For example, many of the gene sets specifying brain cell type markers can be very similar across different brain regions or curated within different databases (e.g., DropViz vs. BrainInABlender vs. Allen Brain Atlas), especially for non-neuronal cell types. It is always reassuring to see some redundancy within results, but there may be questionable added benefit to having the full 766 brain cell type gene sets derived from the Allen Brain Atlas available on Enrichr.

Avoiding excessive redundancy can be particularly important if it turns out that one of those varieties of gene sets (e.g., oligodendrocyte related gene sets) is particularly enriched with differential expression, as the false discovery rate (FDR) corrections performed within many analysis pipelines are highly sensitive to p-value distributions within the results, such that gene set enrichment results that contain a large number of gene sets with low p-values are subjected to a less strict multiple comparisons correction. This issue can be at least partially alleviated by summarizing gene set enrichment results using clustering-based methods, but constructing a custom gene set database with minimal redundancy helps prevent the issue from the start.

##### Gene set quality

Another important consideration when adding or replacing gene sets in Brain.GMT is gene set quality. As discussed above, gene sets derived from low-powered individual differential expression studies are more likely to reflect technical artifacts, therefore, moving forward, we may emphasize the extraction of gene sets from higher powered studies and meta-analyses.

Similarly, for this reason we caution against using gene sets generated by the automated reanalysis of public datasets (e.g., GEO2Enrichr [50]) because of the lack of control for prevalent batch confounds and technical artifacts.

##### Using custom gene sets to run formal comparisons with the published literature

Finally, we found that one of the benefits of using a custom .gmt file like Brain.GMT was the ability to easily and quickly run formal comparisons with the results of similar differential expression studies in the published literature. That said, because of this relative ease, when adding gene sets to Brain.GMT for the purpose of running formal comparisons with the published literature it is particularly important to make decisions about the construction and addition of these gene sets (*i.e*., inclusion criteria) before seeing the results of the gene set enrichment analysis. If decisions about which gene sets to include, how the gene sets are extracted from their respective publications, and the statistical thresholds used to define the gene sets are tailored to produce “the most interpretable” results following an initial analysis this will inflate the likelihood of false discovery, similar to any other form of p-hacking. Likewise, if the decision as to which published studies are used as comparison is made following reading the results of those studies and assessing their similarity to the results of the investigator running the analysis, that will also distort the gene set enrichment analysis in a manner inflating false discovery.

#### Future Development and Remaining Questions

We encourage potential users to reach out to us with any remaining questions or suggestions. We will continue to develop Brain.GMT to enhance interpretation of our own differential expression results. As additions or changes are made, they will be documented on our Github site (*https://github.com/hagenaue/Brain_GMT*).

#### Ethics statements

The full methods used to produce the transcriptional profiling results referenced in our paper are described in detail in their respective publications [14–16] and complied with the National Institutes of Health guide for the care and use of laboratory animals (NIH Publications No. 8023, revised 1978).

#### CRediT author statement

##### Megan H. Hagenauer

Conceptualization, Methodology, Software, Validation, Formal Analysis, Data Curation, Writing – Original Draft, Writing – Review & Editing, Visualization, Supervision, Project Administration

##### Yusra Sannah

Validation, Writing – Original Draft, Writing – Review & Editing, Investigation, Formal Analysis, Software

##### Elaine K. Hebda-Bauer

Validation, Writing – Review & Editing, Investigation, Formal Analysis, Software

##### Cosette Rhoads

Validation, Writing – Review & Editing, Investigation, Formal Analysis, Software

##### Angela M. O’Connor

Validation, Writing – Review & Editing, Investigation

##### Stanley J. Watson, Jr

Funding Acquisition, Resources

##### Huda Akil

Funding Acquisition, Resources, Writing – Review & Editing

## Acknowledgments

This study was supported by the Hope for Depression Research Foundation (HDRF) (HA), NIDA U01DA043098 (HA, JL, AAP), ONR 00014-19-1-2149 (HA), the Pritzker Neuropsychiatric Research Consortium (HA, SJW), Grinnell College Center for Careers, Life, and Service, and the University of Michigan Undergraduate Research Opportunities Program (UROP).

We would also like to thank Dr. Elissa Chesler and Dr. Erich Baker for answering questions about the GeneWeaver database and API, and thank Dr. Paul Pavlidis and Dr. Ogan Mancarci for answering questions about the Gemma database and API.

